# Deep Proteome Profiling of Human Mammary Epithelia at Lineage and Age Resolution

**DOI:** 10.1101/2021.02.02.429276

**Authors:** Stefan Hinz, Antigoni Manousopoulou, Masaru Miyano, Rosalyn W. Sayaman, Kristina Y. Aguilera, Michael E. Todhunter, Jennifer C. Lopez, Lydia L. Sohn, Leo D. Wang, Mark A. LaBarge

**Author notes:** Authors contributed equally (Stefan Hinz, Antigoni Manousopoulou). Corresponding author: Mark LaBarge.

## Abstract

Age is the major risk factor in most carcinomas, yet little is known about how proteomes change with age in any human epithelium. We present comprehensive proteomes comprised of >9,000 total proteins, and >15,000 phosphopeptides, from normal primary human mammary epithelia at lineage resolution from ten women ranging in age from 19 to 68. Data were quality controlled, and results were biologically validated with cell-based assays. Age-dependent protein signatures were identified using differential expression analyses and weighted protein co-expression network analyses. Up-regulation of basal markers in luminal cells, including KRT14 and AXL, were a prominent consequence of aging. PEAK1 was identified as an age-dependent signaling kinase in luminal cells, which revealed a potential age-dependent vulnerability for targeted ablation. Correlation analyses between transcriptome and proteome revealed age-associated loss of proteostasis regulation. Protein expression and phosphorylation changes in the aging breast epithelium identify potential therapeutic targets for reducing breast cancer susceptibility.

## INTRODUCTION

Aging is a major risk factor for most diseases in humans, and it is the greatest risk factor for solid tissue carcinomas. Aging increases susceptibility to cancer initiation, but the underlying reasons for this are unclear (Marusyk and DeGregori, 2008). A majority of effort spent characterizing effects of age in various human tissues have utilized transcriptome and genome-wide analyses of whole tissues (Berchtold et al., 2008; de Vries et al., 2017; Glass et al., 2013; Haakensen et al., 2011). There is poor correspondence between transcriptome and proteome (Kelmer Sacramento et al., 2020; Wang et al., 2019), and proteomic approaches to investigate effects from aging in human tissues have been limited to whole tissues, obscuring lineage-specific changes (Johnson et al., 2020).

We are using breast as a model for understanding how aging increases susceptibility to cancer because cell lineages are well-defined with biomarkers, and surgically discarded normal tissue is obtainable from cosmetic procedures. Age is the greatest risk factor for breast cancer as over 75% of new breast cancer diagnoses are made in women aged 50 or older (Benz, 2008). Whole breast tissue analyses identified a number of directional changes in gene expression (Lee and Lee, 2017; Song et al., 2017). However, aging is associated with significant shifts in proportions of epithelial and stromal cells so whole tissue signals mask age-dependent intrinsic changes to the different cell types (Garbe et al., 2012). Proteomic examination of breast-related tissue has been mostly limited to breast cancer cell lines, which revealed pathways that are active in different cancer subtypes but teaches nothing about aging or cancer susceptibility (Kalocsay et al., 2020; Lawrence et al., 2015). Indeed, there is a paucity of proteomic resources for understanding the effects of aging in breast, or in any other human tissue. We performed proteomics on purified populations of the two principle epithelial lineages in breast: the contractile and tumor suppressive myoepithelial cells, and the secretory luminal epithelial cells, which are thought to be the cancer cells of origin for the luminal subtypes of breast cancer that are most age-related (Prat and Perou, 2011).

Here, we examined age-dependent changes in the proteomes and phosphoproteomes of human mammary luminal epithelial and myoepithelial cells from 10 women who were aged 19-68 years at the time of surgery. We identified differentially expressed proteins, enriched gene sets, weighted protein correlation network modules that change with age, and examined the age-dependent loss of matched mRNA-protein expression patterns. Previously identified age-dependent increases in expression of basal genes in luminal cells with age were verified proteomically, and extended to include the receptor tyrosine kinase AXL and other proteins that control epithelial plasticity and epithelial to mesenchymal transitions. Heretofore unrecognized changes in the DDR1/PEAK1 pathway in luminal cells were identified as a potential age-dependent vulnerability, and we performed proof of principle drug-targeted ablation of older luminal epithelial cells. Correlation analyses of genome-wide transcriptome and proteome data with subsequent biological validation showed age-dependent loss of proteostasis in luminal cells. Our data presents novel opportunities to understand the cellular aging processes at the proteomic and phospho-peptide levels in two epithelial lineages from a normal human tissue.

## RESULTS

### Deep proteomic and phosphopeptide quantification of the mammary epithelium

To investigate age-dependent changes in the mammary epithelium, we utilized tandem mass tag mass spectrometry (TMT-MS) to quantify the proteomic and phosphopeptide landscape of normal human mammary epithelial cells (HMECs) from 5 younger (age < 30 years) and 5 older (age >50 years) women (**Figure 1A**). Lineages were FACS enriched prior to TMT-MS into luminal epithelial (LEp, CD133^+^/CD271^−^) and myoepithelial (MEp, CD133^−^/CD271^+^) cells. The two different lineages from the same specimen were examined in tandem in separate runs. The proteomic analyses yielded >9000 proteins (at 5% spectral mapping false discovery rate (FDR)) for both LEp and MEp lineages, the majority (>84%) were detected in both data sets (**Figure 1B**). Samples were analyzed as all LEp cells in tandem or all MEp cells in tandem enabling within lineage comparisons, but not between lineage comparisons. The phosphopeptide analyses detected 9799 and 15713 distinct phosphopeptides for the LEp and MEp data sets, respectively (**Figure 1B**). The overall distributions of the median log2 abundances were comparable between the datasets (**Figure 1C**). Samples consisting of older women presented wider distributions indicative of a larger range of protein expression levels. A global overview of the LEp and MEp MS datasets demonstrated that the largest number of proteins observed were for proteins and peptides categorized as transcription factors or mitochondria associated proteins (**Figure 1D-G**). Our previous work highlighted KRT14 and KRT19 as exemplar genes and proteins that show age-dependent changes in LEp cells (Garbe et al., 2012; Pelissier Vatter et al., 2018) and we also detected an increase in KRT14 and a decrease of KRT19 expression with age in these MS data (**Figure 1H**). Additionally, we demonstrated examples of age-dependent expression of ZNF542P and AVIL in the MEp cells as a characterization of the changing proteomic composition of the mammary epithelium (**Figure 1I**). These data provided a high-level overview of the produced datasets that vastly enhanced a proteome and phosphopeptide wide understanding of the aging mammary epithelium

**Figure 1:**
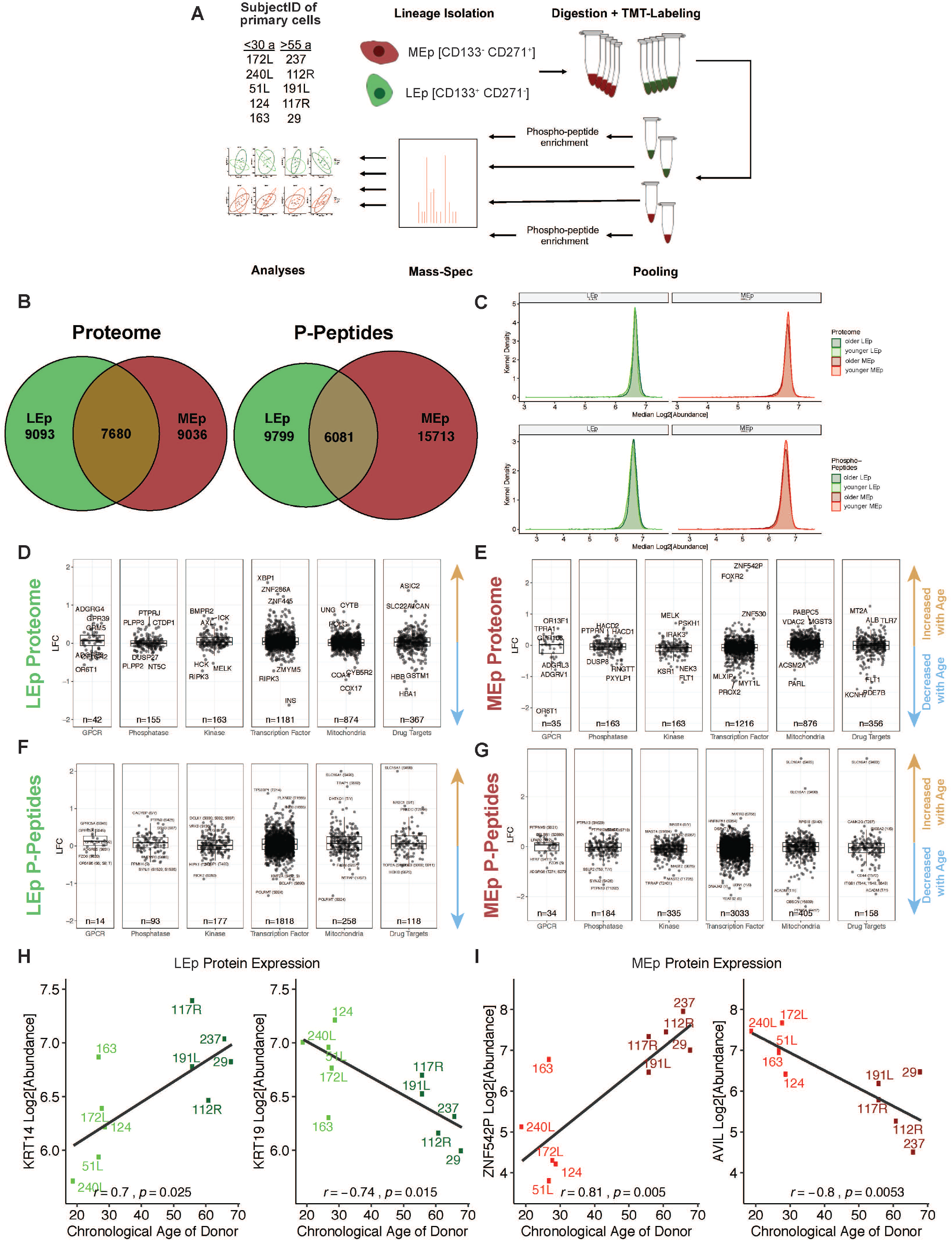
Proteomic and phosphopeptide lineage specific profiling of primary mammary epithelial cells. A) Flowchart of experimental workflow. B) Venn diagram of quantified proteins and phosphopeptides across datasets. C) Density plot of log2 transformed proteins abundances. D-G) Overview of proteins’ and phosphopeptides’ log2 fold changes (old vs young) in selected functional protein classes (young: from women <30 years (n = 5), old: > 50 years old (n = 5)). H) KRT14 and KRT19 expression as a function of age in luminal epithelial (LEp) cells. I) ZNF542P and AVIL expression as a function of age in myoepithelial (MEp) cells.

### Age-dependent changes are more pronounced in luminal epithelial cells

To assess the contribution of age to the proteomic and phosphopeptide landscape we conducted nonlinear dimension reduction analyses using both uniform manifold approximation and projection (UMAP) as well as t-distributed stochastic neighbor embedding (t-SNE) (**Figure 2**). A robust age group-related separation was observed in the tSNE and UMAP plots for LEp cells along with a wider 95% confidence ellipse indicative of increased expression variance in LEp cells with age (**Figure 2A,C**). This observation was especially pronounced in 3-dimensional (3-D) tSNE data. Notably, MEp cells were less clearly separated by age and the 95% confidence ellipses demonstrated a greater overlap for both the proteomic and phosphopeptide data, indicative that effects of aging were less pronounced in these cells (**Figure 2B,D**). These analyses show that the age-dependent changes were more prominent in LEp cells compared to MEp cells.

**Figure 2:**
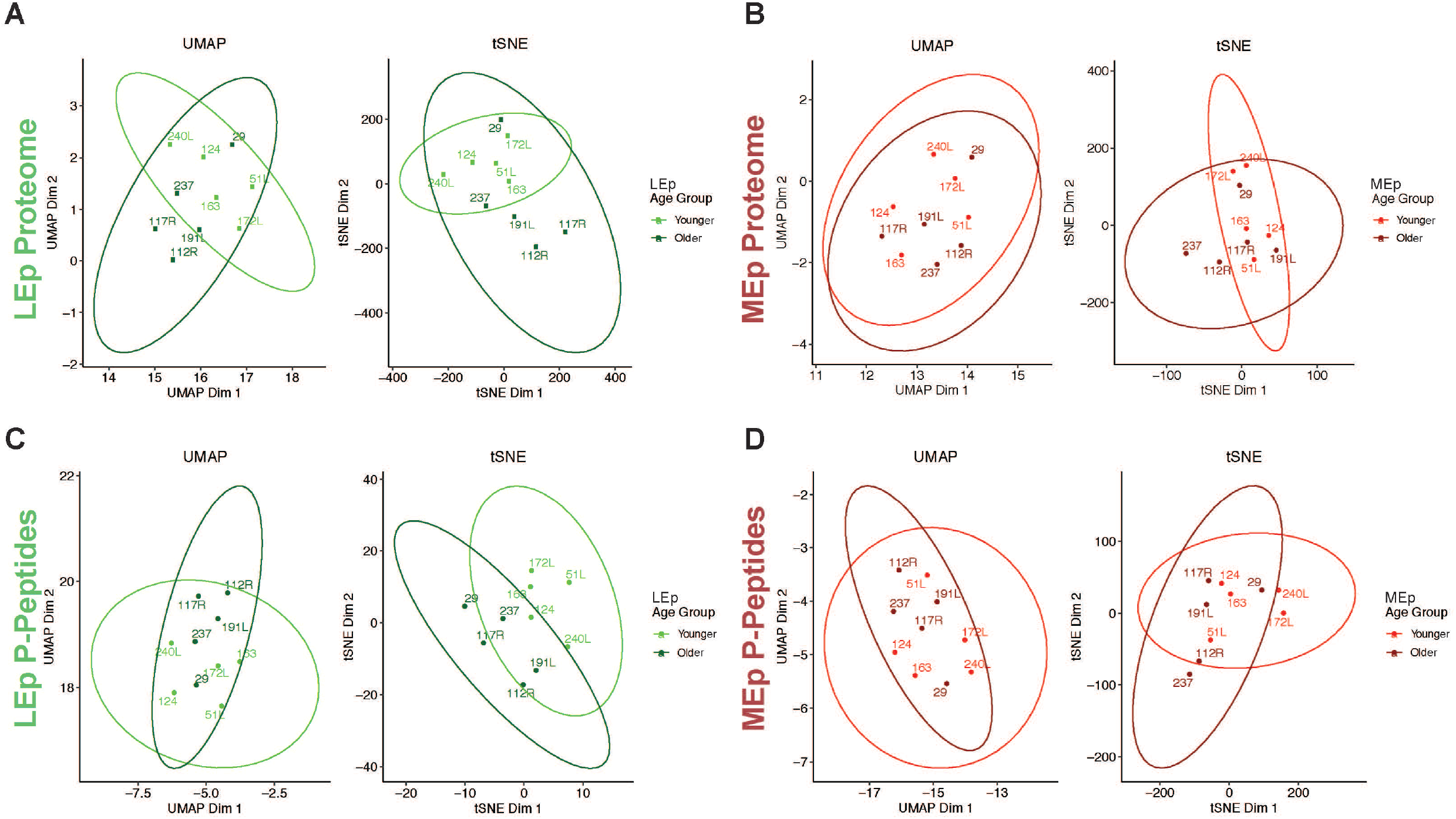
Dimension reduction analyses of datasets. UMAP and tSNE plots with 95% confidence ellipse of HMEC datasets by age (young: from women <30 years (n = 5), old: > 50 years old (n = 5)) for proteins in A) LEp, B) MEp, and phosphopeptides (p-peptides) for C) LEp, D) MEp.

### Aging signatures of the mammary epithelium

Differential expression analyses were conducted to identify proteins that are age-dependent and we detected 155 differentially expressed proteins (DEPs) in LEp cells and 234 in MEp cells (cutoff of adjusted p value < 0.05, only the 100 most significant proteins shown in **Figure 3A-B**). In the LEp cells 114 DEPs were upregulated, including KRT14, KRT10, KRT15, and AXL, while only 41 DEPs were downregulated. Inversely, for MEp cells we detected that 197 of the 234 DEPs were downregulated and 37 were upregulated. Differential expression analyses were also performed for phosphopeptides (DEpP) in LEp cells (**Figure 3C**) and MEp cells (**Figure 3D**) and identified 77 DEpPs in LEp cells with 61 upregulated and 16 downregulated peptides. We identified 365 DEpPs for MEp cells; 53 were upregulated and 312 were downregulated.

**Figure 3:**
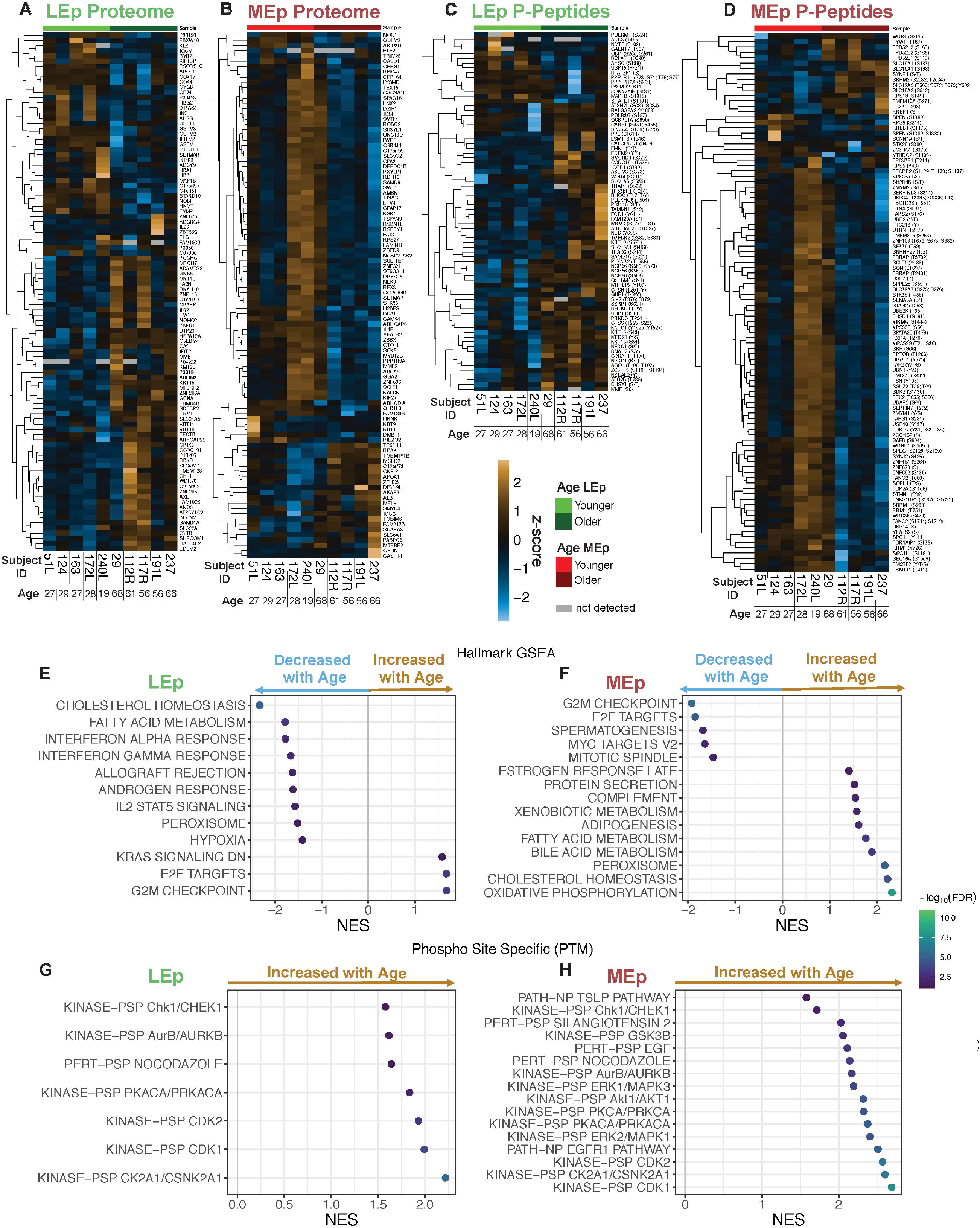
Differential expression analyses between younger and older HMECs by lineage. Heatmaps of the top 100 differentially expressed proteins for LEp (A), MEp (B) and phosphopeptide (p-peptide) level (z score scale). Differentially expressed phosphopeptides for LEp (C) and MEp (D). Gene set enrichment analyses of protein samples for E) LEp and F) MEp cells. Phosphorylation site-specific signature enrichment analysis for G) LEp and H) MEp cells.

Gene set enrichment analyses (GSEA) were conducted for the proteome datasets using a ranked gene list based on the product of log2 fold change (LFC) and the inverse of the p value to identify hallmark signatures enriched with age for LEp cells (**Figure 3E**) and MEp cells (**Figure 3F**). In LEp cells we identified 12 significantly enriched hallmark terms (FDR < 0.05). Three gene sets were enriched in older LEp cells, including reduced KRAS signaling, expression of E2F targets, and G2M checkpoint. In younger LEp cells we identified 9 genemsets enriched including IFN-alpha and -gamma, allograft rejection, IL2 STAT2 signaling, cholesterol homeostasis, fatty acid metabolism, peroxisome, and hypoxia (**Figure 3E**). Of the 15 significantly enriched genesets for MEp cells, 10 genesets are enriched in older MEp cells including oxidative phosphorylation, peroxisome, cholesterol homeostasis, bile acid metabolism, fatty acid metabolism, adipogenesis, xenobiotic metabolism, complement, protein secretion, and late estrogen response. In younger MEp cells 5 gene sets were found to be enriched including G2M checkpoint, targets of the transcription factor E2F, spermatogenesis, MYC targets version 2, and mitotic spindle (**Figure 3F**). For the phosphopeptides a phosphorylation site centric enrichment was performed utilizing PTM signature enrichment analysis (PTM-SEA). In LEp cells 7 significantly (FDR < 0.05) enriched signatures were identified including kinase activity associated with cell cycle control (CHEK1, CDK2, CDK11, CSNK2A1, and PRKACA) (**Figure 3G**). In the phosphoproteome of MEp cells 16 significant (FDR < 0.05) signatures were found to be enriched. These also contained multiple signatures of kinases associated with cell cycle control (CDK1, CSNK2A1, CDK2, PRKCA,AURKB, and CHEK1), and pathway activation of key signaling cascades (AKT1, ERK1, EGFR1, and GSK2B) (**Figure 3H)**. For both LEp and MEp cells these PTM-SEA results are consistent with the protein centric GSEA analysis and increase the understanding of signaling events in the mammary epithelium. Collectively, these analyses show that aging-dependent changes are highly lineage specific. Moreover, for certain gene sets (including cholesterol homeostasis, E2F targets, and G2M checkpoints) the age-dependent regulation was inverse when comparing MEp cells to LEp cells.

### Identification of age-dependent co-expression modules and targetable factors

Weighted correlation network analyses (WGCNA) were performed to identify novel groups of proteins with expression profiles that correlate with age. A co-expression network based on the protein and phosphopeptide expression profiles was constructed. A soft threshold power β of 18, was determined in all datasets to reach a degree of independence over 0.8. Eighteen co-expression modules (fast cluster algorithm with a minimum module size of 100 proteins) were identified in the LEp proteome with 4 of the modules significantly (adjusted p < 0.1) correlated with chronological age (**Figure 4A-B**). Modules “grey60” (r_bicor_ = 0.92, adjusted p = 0.003) and “lightcyan” (r_bicor_ = −0.75, adjusted p = 0.06) were positively correlated with age, while modules “lightgreen” (r_bicor_ = −0.75, adjusted p = 0.06) and “midnightblue” (r_bicor_ = −0.84, adjusted p = 0.02) were negatively correlated with age. The “grey60” module contained 111 proteins (**Figure 4C**), while module “midnightblue” contained 121 proteins. The expression patterns in these modules were homogeneous per age group as indicated by eigengene expression plots (**Figure 4C-4F**). The most interconnected proteins of the “grey60” (positively correlated with age) and “midnightblue” (negatively correlated with age) modules are shown in network plots (**Figure 4G-H**). KRT14 was identified as one of the most interconnected proteins within the “grey60” module, and other key signaling proteins such as PEAK1, IPPK, and CDK13 were also identified to be positively correlated with age (**Figure 4G**). Among the key negatively correlated proteins in the “midnightblue” module were KRT19, ALDH1A3, UBA6, and AVIL (**Figure 4H**).

**Figure 4:**
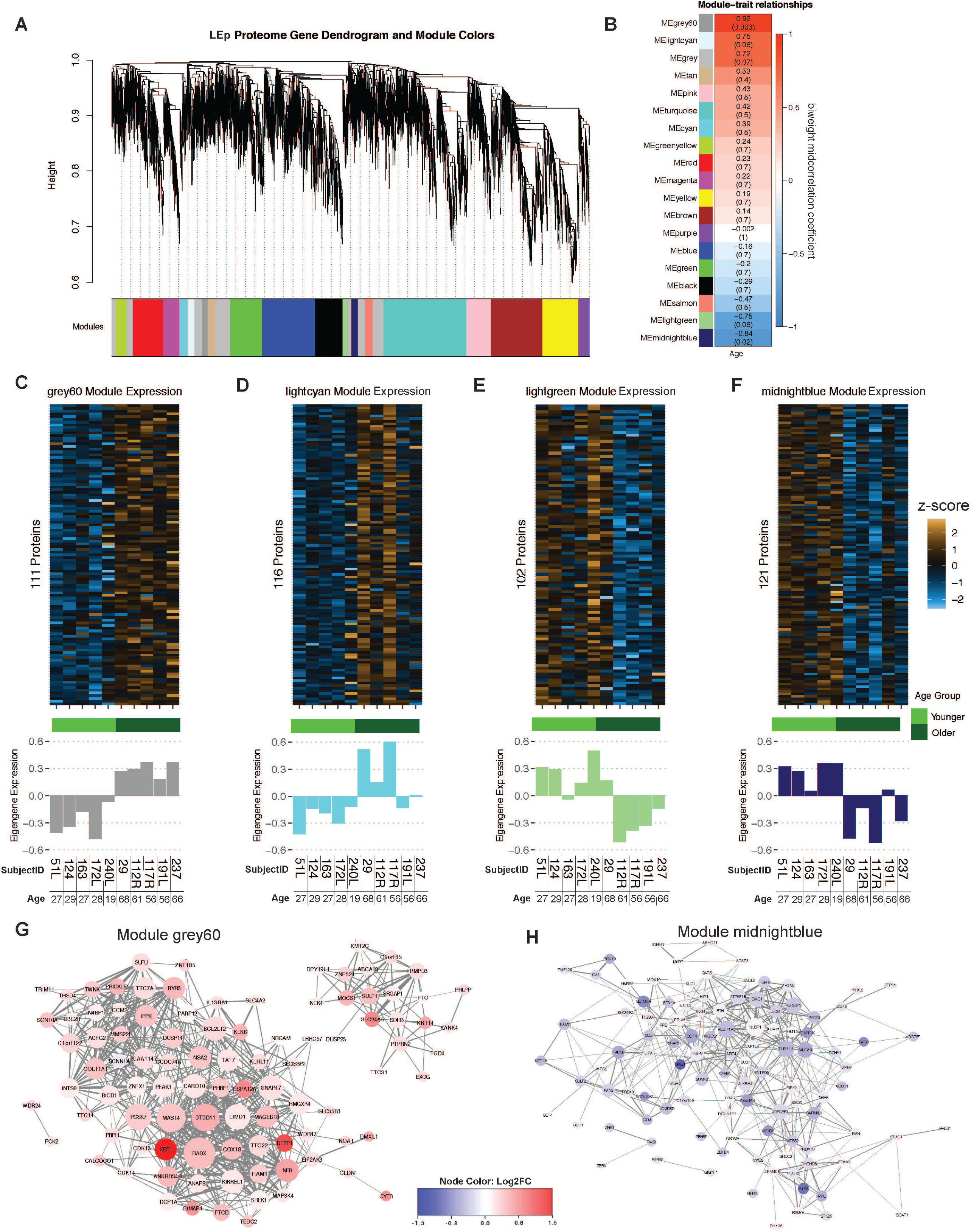
Weighted correlation network analyses for protein expression in LEp cells. A) Protein dendrogram acquired by average linkage hierarchical clustering. Module assignment determined through dynamic tree cut and indicated by color row. B) Module eigengene correlation with age. Heatmap color refer to the biweight midcorrelation and the Benjamini-Hochberg adjusted p-value is given in parenthesis. C-F) Heatmaps of modules significantly correlation with age (z score scale) and barplots of eigengene expression of samples within the modules. Network plots of the most correlated modules G) “grey60” and H) “midnightblue”. Modules are colored by the protein expression log2 fold change and the size of the nodes are relative to the number of connections. The network only displays connections if the topological overlap is above a threshold of 0.08. Nodes with less than 3 connections were removed from plot.

Importantly, the identification of key known age-dependent proteins such as KRT14 and KRT19 within the modules serve as validation of the biological relevance of the analyses. Additionally, we validated PEAK1 as an additional central kinase positively correlated with age in our LEp cells. PEAK1 is heretofore unexamined in the context of the aging breast epithelium and is a factor established downstream of discoidin domain receptor 1 (DDR1), which is involved in pro-tumorigenic signaling in other carcinoma model systems (Aguilera et al., 2017; Hur et al., 2017; Saby et al., 2018). PEAK1 is a downstream effector of DDR1 in pancreatic cancer (Aguilera et al., 2017; Aguilera et al., 2014) and we thought it remarkable to identify its age-dependent expression here. As a biological validation of the WGCNA analysis we examined total PEAK1 protein expression levels as a function of age in HMECs and showed an increased expression with age via western blot (**Figure 5A**). By utilizing the DDR1-specific drug 7rh, which was previously utilized to target DDR1 signaling in pancreatic cancer (Aguilera et al., 2017), phosphorylation of both DDR1 and PEAK1 were reduced in HMECs from older women (**Figure 5B**). Dose-response analyses were used to assess cell viability in different concentrations of 7rh. Sensitivity to 7rh significantly increased with age (2-way ANOVA p < 1e-4, **Figure 5C**).

**Figure 5:**
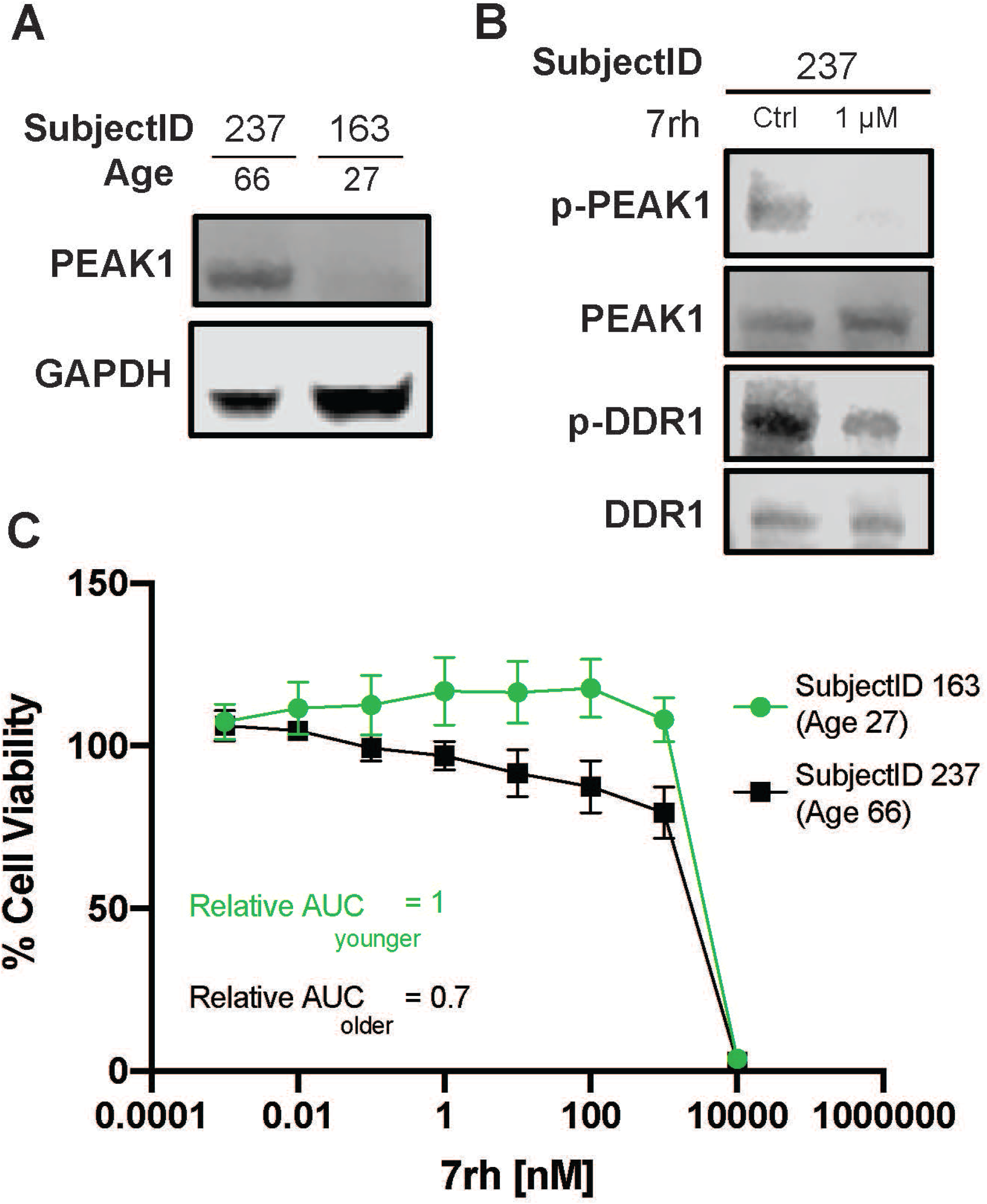
Targeting PEAK1 expression through DDR1 inhibition: A) Western blot of PEAK1 expression in younger (SubjectID 163, age 27) and older (SubjectID 237, age 66). B) Western blot of phospho-PEAK1 and phospho-DDR1 after inhibition of DDR1 mediated signaling through 24h treatment with 1 μM 7rh. C) Cell viability assay (cell titer glo) after 72h treatment with 7rh.

Continuing the WGCNA analyses to interrogate the phosphopeptides, ten modules were identified for the LEp dataset including the “brown” module (r_bicor_ = 0.67, P = 0.03) which included 298 phosphopeptides significantly correlated with age (**Figure 6A-C**). We generated a co-expression network plot that depicts up and down-regulated proteins (**Figure 6D**) including cell cycle related peptides, a YAP pathway associated transcription factor (TEAD3), multiple RAB peptides, and key pathway kinases (MAPK2K7 and ILK). These results are consistent with PTM-SEA (**Figure 3G**). No co-expression modules significantly correlated with age in the MEp datasets, indicating a reduced effect of aging on co-expression networks. Thus, WGCNA analyses identified key protein correlation networks for LEp cells that were associated with age and we presented data that experimentally confirmed the findings and targetability.

**Figure 6:**
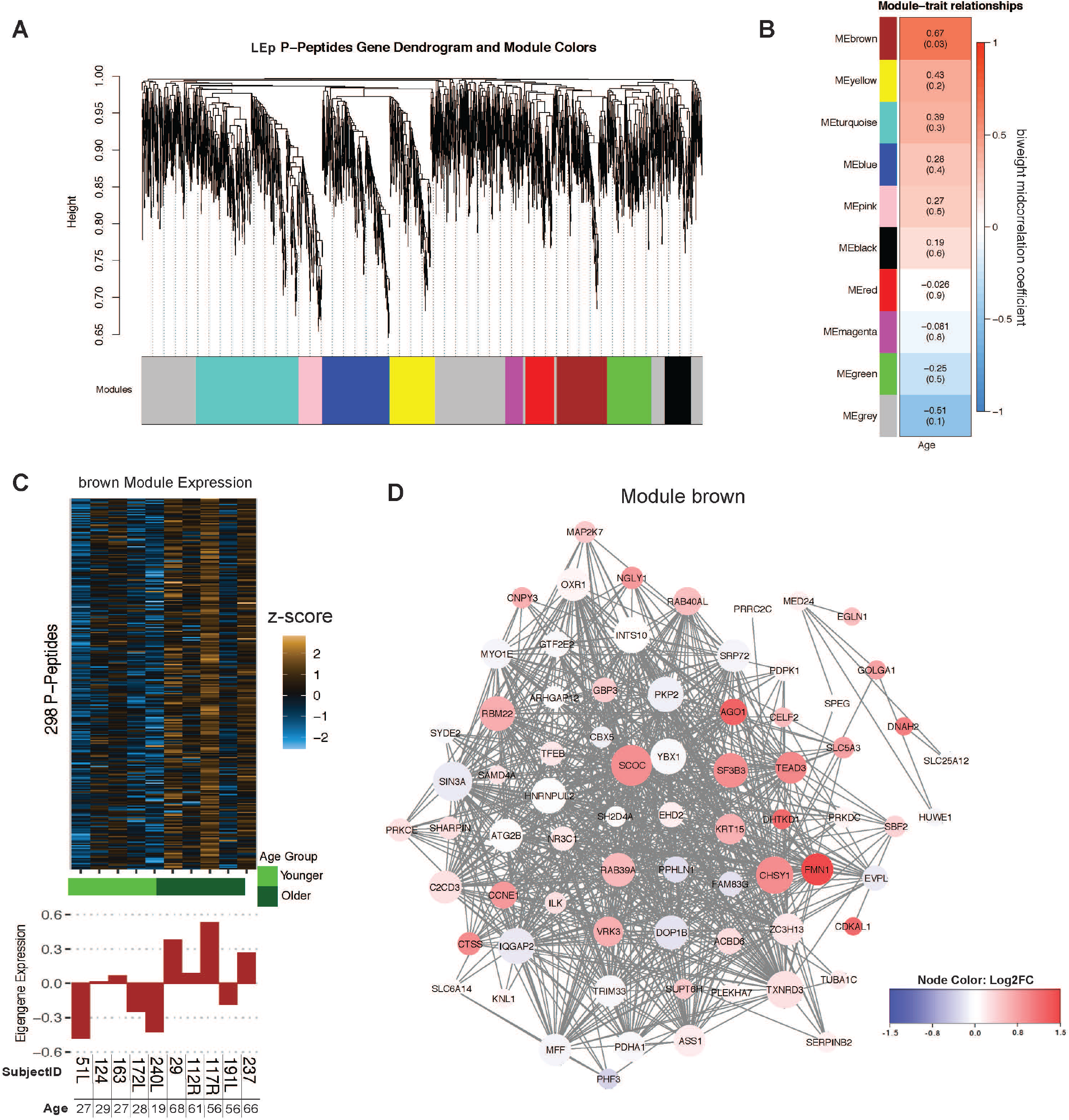
Weighted correlation network analyses for phosphopeptide expression in LEp cells. A) Protein dendrogram acquired by average linkage hierarchical clustering. Module assignment determined through dynamic tree cut and indicated by color row. B) Module eigengene correlation with age. Heatmap color refer to the biweight midcorrelation and the p-value is given in parenthesis. C) Heatmap of brown module (z score scale) and barplots of eigengene expression of samples. D) Network plots of the most correlated modules. Modules are colored by the proteins log2 fold change and the size of the nodes are relative to the number of connections. The network only displays connections if the topological overlap is above a threshold of 0.2. Nodes with less than 3 connections were removed.

### Correlation of transcriptome and proteome

We correlated RNAseq-derived transcriptomes with proteomes (**Figure 7A-B**) and demonstrated an observed median correlation of 0.19 (r_bicor_ = 0.19) for LEp cells and 0.04 (r_bicor_ = 0.04) for MEp cells. The correlation of keratins and ribosomal transcripts and proteins was greater in LEp cells compared to MEp cells. Whereas protein transport genes and proteins showed a lower correlation in LEp cells compared to MEp cells. Using the gene-protein correlation pairs we investigated differential correlation changes by age (**Figure 7C-D**). For LEp cells four transcript-protein pairs that changed the direction of correlation with age were identified. A positive correlation in younger cells that changed to a negative correlation in older cells was observed for ACAT1 (mitochondria associated), PRKACB (serine/threonine protein kinase), and TTC39C (unknown function). A negative correlation in younger strains and strong positive correlation with age was identified for NOP16 (ribosomal protein), which suggested a possible change of the ribosomal complex assembly machinery. In MEp cells five differentially correlated transcript/protein pairs were detected. POLR2M (RNA Pol II Subunit M), PPP2CB (protein phosphatase), and PEX6 (peroxisomal protein import) were highly positively correlated in younger strains and anticorrelated with age. Two proteins/transcripts that gained correlation with age were PPCS (metabolomic protein) and ANKMY1 (ankyrin repeat and MYND domain containing 1).

**Figure 7:**
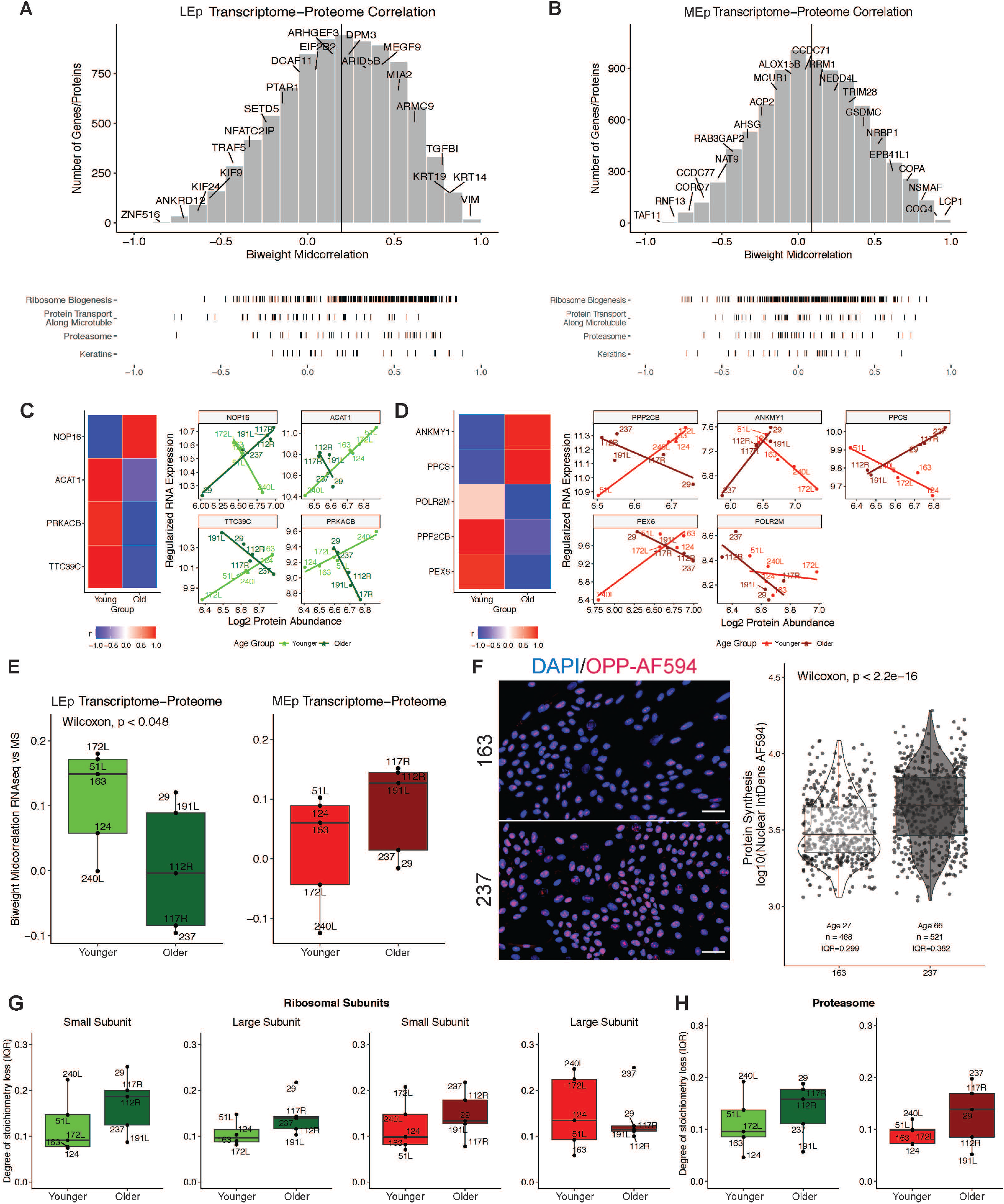
Correlation between protein and RNA expression. Histogram of the biweight midcorrelation distribution between protein and RNA expression for A) LEp and B) MEp with ridgeplot depicting distribution of selected gene groups curated by HGNC. C) Differentially correlated transcripts-proteins pairs by age for LEp cells and D) MEp cells. E) Box and whisker plots of protein-transcript biweight midcorrelation by age and lineage (Wilcoxon one sided signed rank test). F) Protein synthesis quantification using OPP-AF594 with quantification of nuclear integrated density per cell for 2 strains (scale bar 50 μm). Change in protein expression stoichiometry of G) ribosomal and H) proteasomal proteins by age and lineage.

We further investigated the correlation between gene expression and protein levels with age (**Figure 7E**). Whereas MEp cells did not show loss of correlation, there was a significant age-dependent loss of correlation between the transcriptome and proteome in LEp cells (Wilcoxon test one sided, p = 0.048). Since proteostasis is considered a hallmark of aging (Kaushik and Cuervo, 2015; Lopez-Otin et al., 2013), it was investigated whether the loss of correlation was attributable to an alteration of overall protein production. A significant increase in protein production was observed in an older HMEC strain (age 66) as compared to a younger one (age 27) (**Figure 7F**) utilizing a OPP-AF594 based protein synthesis assay. To further investigate the dysregulation of the translational machinery we evaluated stoichiometric changes of the proteasomal and ribosomal subunits. Janssens et al. (2015) showed that loss of stoichiometry (uncoordinated change of expression) can alter the assembly of multiprotein complexes, which ultimately alters protein homeostasis (Kelmer Sacramento et al., 2020). Using the HUGO gene group annotation for the ribosomal subunits and proteasome associated proteins we determined a partial loss of stoichiometry for both LEp cells and MEp cells in an age-dependent manner (**Figure 7G-H**). These data link the loss of the mRNA-protein correlation with age and dysregulation of the proteostasis machinery.

## DISCUSSION

Age is the most important risk factor for developing cancer. Thus, there is an urgent need to investigate the biology and mechanisms underlying this increased susceptibility in order to develop prevention and screening modalities. Here we present the first comprehensive and in-depth characterization of the proteomic and phosphopeptide changes of the normal aging mammary epithelium at lineage resolution. We present differentially expressed proteins and phosphopeptides and have identified key co-expression modules that correlate with the chronological age of the tissue donor. We also utilized matched RNAseq data to investigate the age-dependent decoupling between transcriptome-proteome correlations. Current proteomic studies that investigate human aging predominantly focus on blood samples generally lacking epithelial tissue (Johnson et al., 2020; Ubaida-Mohien et al., 2019). Moreover, proteomic profiling of breast-derived cells involve the use of abnormal and cancer cell lines (Kalocsay et al., 2020; Lawrence et al., 2015). Here we present a data resource for gaining insight into expression and phosphorylation of over 9000 proteins in normal primary breast epithelia at lineage- and age-resolution.

We identified a number of age-dependent protein and phosphopeptide changes that could be leveraged as aging biomarkers or targets in future studies of aging in the breast. The most prominent age-dependent changes were identified in LEp cells, consistent with our previous findings (Pelissier Vatter et al., 2018). Previous studies by our group characterized changes in human mammary epithelia with age and one of the most striking changes between cells from young compared to older HMECs is keratin expression in luminal cells (Garbe et al., 2012; Pelissier Vatter et al., 2018). KRT14 is a lineage specific marker for MEp cells in young women, but LEp cells of older women acquire expression of this intermediate filament. KRT14 downregulation in cancer cell lines showed reduced proliferation, tumorigenicity, and reduced activation of the AKT pathway and present a potential target to reduce cancer susceptibility with age (Alam et al., 2011). Here we have detected a significant increase of KRT14 expression, as well as KRT10 and KRT15, in our LEp dataset and detected KRT14 in the WGCNA module that is most correlated with age. Given that the genes for these keratins, in addition to *KRT19*, are proximally located to one another on chromosome 17 suggests that they may be co-regulated in an age-dependent manner. Taken together, our analyses of TMT-MS data for known changes validates the approach we have taken, giving confidence for future interrogation of other targets presented here.

WGCNA analysis revealed the age-dependent expression of PEAK1 (pseudopodium enriched atypical kinase 1), a signaling kinase previously unassociated with aging. PEAK1 is a non-receptor tyrosine kinase ubiquitously expressed and dysregulated in several cancer models including the pancreas (Aguilera et al., 2017; Aguilera et al., 2014) and breast (Wang et al., 2010). Aguilera et al. (2014) established PEAK1 as a downstream target of DDR1 (discoidin domain receptor 1) that can be pharmacologically targeted with a small molecular inhibitor, 7rh (Aguilera et al., 2017; Gao et al., 2013). We show sensitivity to 7rh in an age-dependent manner and confirm decreased phosphorylation of PEAK1 after treatment. These experiments suggest a role of PEAK1 in the mammary epithelium that is age-dependent, targetable, and could lead to novel breast cancer prevention intervention strategies that include prophylactic ablation of LEp cells or their precursors that exhibit increased PEAK1 expression and activity. Mature LEps are thought to be the cells of origin for the luminal subtype breast cancers (Prat and Perou, 2010), 80% of which are age-associated, and DDR1/PEAK1 represents a heretofore unknown vulnerability.

Our findings uncover a loss of transcriptome-proteome quality control in an age-dependent manner that is most prominent in LEp cells. We observed an increased overall protein production and reduced correlation to the transcriptome. Loss of proteostasis control has previously been proposed as a hallmark of aging (Balch et al., 2008) and utilizing the approach proposed by Janssens et al. (2015) we established an increase of interquartile range of multiple protein complexes involved in proteostasis. Dysregulation of the stoichiometry of these complexes could contribute to the decoupling that has been extensively studied in aging models of yeast and killifish (Janssens et al., 2015; Kelmer Sacramento et al., 2020). We speculate that this decoupling may be the result, or the cause, of age-dependent increased transcriptional variance that has been reported in immune, pancreatic, and breast cells (Enge et al., 2017; Martinez-Jimenez et al., 2017). The loss of correlation between RNA and protein highlights the need for protein evidence-based investigations, for example pediatric cancer subtypes were identified based on only proteome data (Petralia et al., 2020).

The key to this study is the utilization of pre-stasis epithelial cells from reduction mammoplasty tissues to examine the aging process, a model system which we generated and is well established (Garbe et al., 2012; Garbe et al., 2014; Labarge et al., 2013; Pelissier et al., 2014; Pelissier Vatter et al., 2018). In our present study, HMECs were isolated from 10 different reduction mammoplasties spanning an expansive age range from 19 to 68 years. LEp cells and MEp cells were isolated from finite primary cells that were not exposed to immortalization factors, do not possess gross genetic alterations, transformations, nor genomic instabilities (Stampfer et al., 2013). This is unlike the cell lines, such as MCF10A or HMLER, which display high genetic instability and often require overexpression of transformation factors, converting these cell lines into an abnormal cell state (Stampfer et al., 2013). The power of using systems like these is that, unlike primary tissues which can usually only support n of 1 analysis or functional assays, this HMEC system enables one to follow-up and test predictions from the proteomics data in cell-based assays.

The resource we provide here is a remarkable dataset that allows in-depth analyses of aging in normal human mammary epithelium. We observe large scale expression changes especially in the luminal subpopulation of the epithelium, which is the most likely culprit of most age-associated breast cancers. Ultimately, these data could lead to further interrogations that may uncover novel aging biomarkers, high risk identifiers, and therapeutic interventions to prevent or treat age-associated breast cancers.

## METHODS

### EXPERIMENTAL MODEL AND SUBJECT DETAILS

#### Cell Culture

Primary HMECs at passage 4 were grown at 37°C in M87A medium containing cholera toxin and oxytocin at 0.5 ng/ml and 0.1 nM (Garbe et al., 2009). HMEC strains used in this study were 51L, 124, 163, 172L, 240L, 29, 112R, 117R, 191L, and 237. Media was changed every 48h with the last media change at 24h prior to cell dissociation.

#### Flow Cytometry

Cells dissociated from primary HMEC strains (passage 4) were stained with anti-human CD271-PerCP/Cy5.5 (Biolegend #345122) and anti-human CD133-PE (Biolegend #372804) by following standard flow cytometry protocol. Cells were sorted by S3 Cell Sorter (Bio-Rad). After sorting, cells were washed three times with PBS, snap frozen, and stored at −80°C.

#### Mass Spectrometry

Cell pellets were dissolved in 0.5 M triethylammonium bicarbonate (TEAB) (T7408, Sigma-Aldrich, St. Louis, MO, USA) and 0.05% sodium dodecyl sulphate (SDS) (71736, 50 μL in 10mL water/TEAB solution, Sigma-Aldrich, St. Louis, MO, USA), and lysed using pulsed probe sonication (Misonix, Farmingdale, NY, USA). Lysates were centrifuged (16,000 g, 10 min, 4°C) and supernatants were transferred to fresh tubes. Each sample was measured for protein content using the PierceTM BCA protein assay kit-reducing agent compatible per manufacturer’s instructions (23250, Thermo Fisher Scientific, Waltham, MA, US). 100 ug of protein was used per sample, adjusted to the highest volume using lysis buffer (0.5M TEAB, 0.05% SDS). Proteins were then reduced [4 μL of 100 mM tris (2-carboxyethyl) phosphine (TCEP); 646547, Sigma-Aldrich, St. Louis, MO, USA], alkylated [2 μL of 100 mM S-methyl methanethiosulfonate (MMTS); 64306, Sigma-Aldrich, St. Louis, MO, USA] and enzymatically proteolysed using trypsin/LysC (1:25 enzyme:protein ratio; V5072, Promega, Madison, WI, USA). Peptides from each sample were labelled using the ten-plex TMT reagent kit (90110, Thermo Fisher Scientific, Waltham, MA, US). Two ten-plex experiments were performed, one for MEp cells and one for LEp cells from ten subjects. Labelled peptides per ten-plex experiment were mixed and phospho-enrichment was performed using the high-selectTM SMOAC protocol per manufacturer’s instructions (A32992 and A32993, Thermo Fisher Scientific, Waltham, MA, US). The flow-through containing native peptides was offline fractionated using alkaline C4 reverse phase chromatography (Kromasil^®^ C4 HPLC column, 100 Å pore size, 3.5 μm particle size, length × I.D 150 × 2.1 mm, K08670362, Sigma-Aldrich, St. Louis, MO, USA) and each collected fraction was analyzed using the Orbitrap Fusion mass spectrometry system (Thermo Fisher Scientific, Waltham, MA, US).

Unprocessed raw files were submitted to Proteome Discoverer 2.3.0.523 for target decoy search using Byonic. The UniProtKB homo sapiens database (release date Dec 2019) was utilized. The search allowed for up to two missed cleavages, a precursor mass tolerance of 10 ppm, a minimum peptide length of six and a maximum of two dynamic modifications of; oxidation (M), deamidation (N, Q), or phosphorylation (S, T, Y). Methylthio (C) and TMT (K, N-terminus) were set as static modifications. FDR corrected p-value at the peptide level was set at < 0.05 for native proteins and at < 0.01 for phosphopeptides. Percent co-isolation excluding peptides from quantitation was set at 50.

#### Differential Expression

Differential expression was determined utilizing multiple t-tests (1 per protein/phosphopeptide) with pooled standard deviation based on the log2 transformed abundance values. To correct for multiple comparisons the Benjamini, Krieger & Yekutieli (2006) method was deployed and proteins were considered differentially expressed for FDR_BKY_ < 0.05.

#### Gene Set Enrichment Analyses

GSEA were conducted using the fgsea R package (Sergushichev, 2016). Proteome wide expression profiles were ranked using the product of the log2 fold change and inverse of the p-value. Gene sets were accessed through MySigDB V7.0 (Subramanian et al., 2005) and PTM signature enrichment analysis signatures were retrieved via PTMsigDB v1.9.0 (Krug et al., 2019). Terms were considered significantly enriched if FDR < 0.05.

#### Weighted Correlation Network Analysis (WGCNA)

To perform weighted correlation network analysis the R package WGCNA (Langfelder and Horvath, 2008) utilizing the biweight midcorrelation was used. Scale independence and mean connectivity were then tested using a gradient method. Minimum scale independence of at least 0.80 was met for every dataset with soft threshold of 18. The modules were detected by hierarchical average linkage clustering analysis for the protein dendrogram of the topology overlap matrix. To identify age-dependent modules the module-age relationships were calculated using bicor function and significance was determined (FDR < 0.1). Most interconnected proteins and phosphopeptide networks were illustrated using cytoscape 3.8.0 (Shannon et al., 2003).

#### RNA-Protein Correlation Analysis

Correlations between RNA and proteins were calculated using biweight midcorrelation. Differential correlation has been assessed using the fisher r-z-transformation based in the correlation coefficient. P-value was calculated using two-sided t-test and p-value adjustments was performed using Benjamini, Krieger & Yekutieli (2006) method. Changes were considered significant for FDR_BKY_ < 0.1.

#### Protein Synthesis Assay

HMECs were cultured as previously described on 4-well chamber slides. When cells reached subconfluence media was changed. Protein synthesis assay (Thermo Fisher # C10457) was performed according to manufacturer’s instructions. Images were captured in the same imaging session using Nikon DS-Qi2 camera (3s exposure) on a Nikon Ti2 Microscope. Single cell level of nuclear fluorescence was quantified using Cell Profiler 3.1.5.

#### Cell Viability assay

HMECs were plated in white walled 96-well plates at 1500 cells per well. After 24h cells were treated with increasing concentrations of 7rh for 72h. Cell viability was determined using CellTiter-Glo (Promega) and measured using Cytation 3 Cell Imaging Multi-Mode Reader (BioTek). Graphs were plotted and significance was assessed using Graphpad Prism.

#### Western blot

Sub-confluent HMECs were lysed, centrifugated at 13,000 rpm, and supernatend protein concentration was measured using BCA assay. Equal amounts of total protein were separated by SDS-PAGE and transferredonto PVDF membranes. Membranes were underwent blockade for 1 hour in 5% milk in TBS-T. Primary antibody was incubated overnight at 4°C. Membranes were incubated with corresponding HRP-conjugated secondary antibody for 2 hours. Bands were detected using the enhanced chemiluminescence reagent using Oddyssey Fc (Licor).

#### Statistical Analysis

All performed tests were two-sided unless otherwise specified and calculated in R (V3.6.1) or Prism (Graphpad, V9).

### KEY RESOURCES TABLE

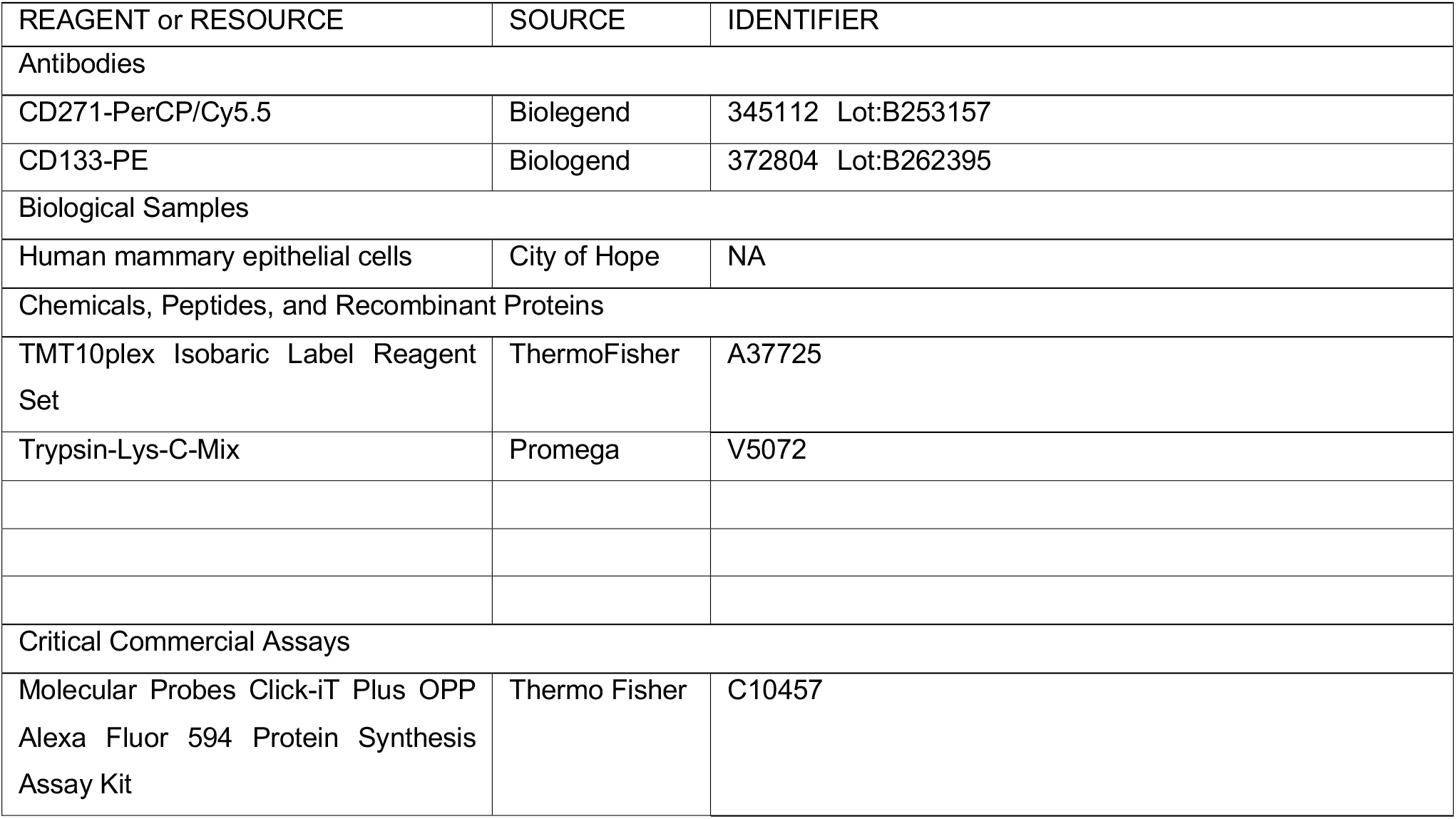
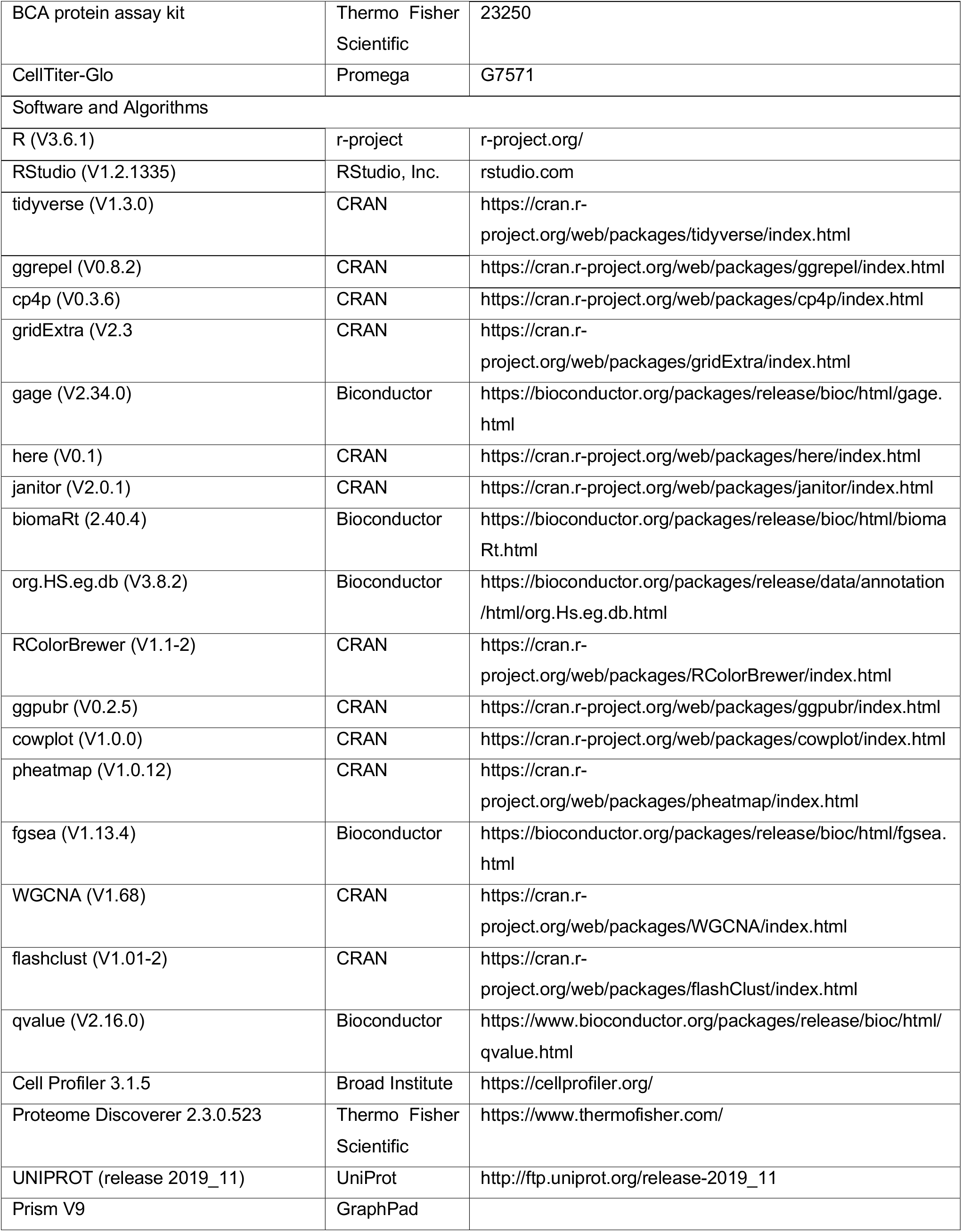

## ACKNOWLEDGEMENTS

We gratefully acknowledge Dr. Rolf A. Brekken for kindly providing 7rh, and our patient advocates Susan Samson and Sandy Preto or providing much needed context. Funding: DOD CDMRP (BC141351 Era of Hope Scholar Award & BC181737), Hilton-Ludwig Foundation, and City of Hope Center for Cancer and Aging to ML; Margaret Early Memorial Research Trust, Pediatric Cancer Research Foundation to LDW; National Institutes of Health/National Cancer Institute (NIH/NCI) grants R01CA237602, U01CA244109, R33AG059206 to ML; and R01EB024989 to LLS and MAL; K08 CA201591 to LDW; NCI Cancer Metabolism Training Program Postdoctoral Fellowship T32CA221709 to RWS; American Cancer Society Postdoctoral Fellowship (131311-PF-18-188-01-TBG) to M.E.T. Research reported in this publication included work performed in the Mass Spectrometry and Proteomics Core, Analytical Cytometry Core, and Integrative Genomics and Bioinformatics Core supported by the National Cancer Institute of the National Institutes of Health under grant number P30CA033572. The content is solely the responsibility of the authors and does not necessarily represent the official views of the National Institutes of Health.

## AUTHOR CONTRIBUTIONS

Conceptualization, SH, AM, MM, KYA, LDW and MAL. Writing - Original Draft, SH, KYA and MAL. Writing – Review & Editing, SH, AM, MM, RWS, KYA, MET, JCL, SDG, LLS, LDW, and MAL. Investigation, SH, AM, MM, KYA, RWS, and JCL. Formal Analysis, SH and AM. Data Curation, SH, SD and AM. Visualization, SH. Supervision, LLS, LDW, and MAL. Funding Acquisition, LLS, LDW, and MAL.

